# *Tripsacum de novo* transcriptome assemblies reveal parallel gene evolution with maize after ancient polyploidy

**DOI:** 10.1101/267682

**Authors:** Christine M. Gault, Karl A. Kremling, Edward S. Buckler

## Abstract

Plant genomes reduce in size following a whole genome duplication event, and one gene in a duplicate gene pair can lose function in absence of selective pressure to maintain duplicate gene copies. Maize and its sister genus, *Tripsacum*, share a genome duplication event that occurred 5 to 26 million years ago. Because few genomic resources for *Tripsacum* exist, it is unknown whether *Tripsacum* grasses and maize have maintained a similar set of genes under purifying selection. Here we present high quality *de novo* transcriptome assemblies for two species: *Tripsacum dactyloides* and *Tripsacum floridanum*. Genes with experimental protein evidence in maize were good candidates for genes under purifying selection in both genera because pseudogenes by definition do not produce protein. We tested whether 15,160 maize genes with protein evidence are resisting gene loss and whether their *Tripsacum* homologs are also resisting gene loss. Protein-encoding maize transcripts and their *Tripsacum* homologs have higher GC content, higher gene expression levels, and more conserved expression levels than putatively untranslated maize transcripts and their *Tripsacum* homologs. These results indicate that gene loss is occurring in a similar fashion in both genera after a shared ancient polyploidy event. The *Tripsacum* transcriptome assemblies provide a high quality genomic resource that can provide insight into the evolution of maize, an highly valuable crop worldwide.

**Core ideas:**

- Maize genes with protein evidence have higher expression and GC content
- *Tripsacum* homologs of maize genes exhibit the same trends as in maize
- Maize proteome genes have more highly correlated gene expression with *Tripsacum*
- Expression dominance for homeologs occurs similarly between maize and *Tripsacum*
- A similar set of genes may be decaying into pseudogenes in maize and *Tripsacum*

---

Developing genomic resources for wild relatives of crops can aid crop breeding because they often possess novel desirable traits, such as resistance to biotic and abiotic stresses. Wild relatives have not passed through a domestication bottleneck and possess untapped genetic diversity. The perennial grass genus *Tripsacum* is the closest sister genus to *Zea* (Bomblies and Doebley, 2005; Mathews et al., 2002), the genus that contains maize (*Zea mays* (L.) ssp. *mays*). As some of the most ecologically dominant grasses in the Americas, *Tripsacum* grasses possess freezing tolerance and perenniality, which are traits that maize lacks. The natural habitat of the *Tripsacum* genus is vast, extending from the northern United States at 42°N latitude to Argentina at 24°S latitude and encompassing both tropical regions and temperate regions with harsh winters (deWet et al., 1982). *Tripsacum* grasses overwinter in sub-zero temperatures in temperate zones. In contrast, freezing temperatures cause maize leaf damage, yield reduction, or death (Li et al., 2016; Carter, 1995; Elmore and Doupnick, 1995). In addition to abiotic stress tolerance, *Tripsacum dactyloides* (L.) L., also known as Eastern Gamagrass, has resistance to three maize pests: the *Striga hermonthica* parasitic weed (Gurney et al., 2003), western corn rootworm (Branson, 1971; Moellenbeck et al., 1995), and *Puccinia sorghi* (common rust) (Bergquist, 1981). The genetic architecture of these beneficial traits has not been fully explored because there are few *Tripsacum* genomic resources. Currently available *Tripsacum* resources include whole genome skim sequencing datasets for diploid and tetraploid *Tripsacum* lines (Chia et al., 2012; Zhu et al., 2016), but an assembled *Tripsacum* genome has not been published. Additionally, 24,616 *Tripsacum dactyloides* isoforms have been assembled and analyzed, revealing that phospholipid biosynthesis genes show rapid evolution in *Tripsacum dactyloides* and may have aided temperate adaptation (Yan, 2017). Maize, the most productive cereal crop in the world (FAOSTAT, http://faostat.fao.org), could be improved by bringing in valuable traits from its wild relative, *Tripsacum*.

The *Tripsacum* gene complement can frame maize gene evolution in a larger context. The *Tripsacum* genus diverged from the *Zea* genus fewer than 1.2 million years ago, which was before maize domestication (Ross-Ibarra et al., 2009). *Tripsacum* and *Zea* make up the subtribe Tripsacinae in tribe Andropogoneae, subfamily Panicoideae, family Poaceae (Soreng et al., 2015). The two genera share a genome duplication event that occurred 5 to 26 million years ago, around the time their shared lineage diverged from the sorghum lineage (Wang et al., 2015; Swigonova et al., 2004; Whitkus et al., 1992; Berhan et al., 1993). Immediately after the tetraploidization event, the ancestor of *Tripsacum* and maize had twenty chromosomes that eventually became rearranged through chromosomal breakage and fusion (Wei et al., 2007; Murat et al., 2010). The maize genome has been reorganized into ten chromosomes whereas *Tripsacum* species have a base chromosome number of 18 and can be diploid (2n=36), triploid (2n=54), or tetraploid (2n=72) (deWet et al., 1982). Despite the difference in chromosome number, there are no evident large-scale chromosomal deletions between the two genera, and they have very similar gene content (Chia et al., 2012).

The maize and *Tripsacum* genomes are still in flux after the whole genome duplication event. Duplicate genes arising from this ancient tetrapolyploidy event are called homeologous genes and can have many evolutionary fates. Most commonly, one homeolog loses gene function by pseudogenization or deletion in absence of selective pressure to maintain both homeologs (Lynch and Conery, 2000; Maere et al., 2005). In other cases, both homeologs are retained after polyploidization. Genes with dosage balance sensitivity tend to maintain their pre-polyploidy stoichiometric ratio and both homeologs are retained (Tasdighian et al., 2017; Conant et al., 2014). Retained homeologs can undergo subfunctionalization or neofunctionalization (Hughes et al., 2014; Pophaly et al., 2015).

Biased gene fractionation (gene loss) occurs when one homeologous region loses more genes than the other homeologous region (Freeling, 2009). In *Arabidopsis*, 85% of homeologous regions showed biased gene fractionation, where one region was 1.6 times more likely to lose genes than its homeolog (Thomas et al., 2006). In maize, 68% of homeologous regions showed biased gene fractionation, where one region was 2.3 times more likely to lose genes than its homeolog (Woodhouse et al., 2010). Genome dominance occurs when the subgenome contributed by one parent retains more genes and has higher gene expression than the subgenome contributed by the other parent (Woodhouse et al., 2014). Maize homeologous regions were partitioned into the dominant subgenome A and the recessive subgenome B based on these gene retention and expression criteria (Schnable et al., 2011). Of the homeologous gene pairs that were retained in maize inbred line B73, more genes were lost from subgenome B than subgenome A in multiple maize inbred lines (Schnable et al., 2011, Brohammer et al. 2018).

The factors controlling genome dominance are not fully elucidated. Garsmeur et al. (2014) postulate that genome dominance may be more likely to occur in species that have experienced allopolyploidy rather than autopolyploidy. Zhao et al. (2017) refine this model and propose that genome dominance occurs when the subgenomes are highly diverged but not when the subgenomes are genetically similar. Supporting this proposal, no subgenome dominance was observed between the closely related soybean progenitor genomes, but subgenome dominance was observed between the divergent maize progenitor genomes (Zhao et al., 2017). Species arising from the same whole genome duplication event behave similarly in the rate of duplicate gene loss and biased fractionation (Sankoff et al., 2010), but any parallels in maize and *Tripsacum* evolution have yet to be studied. Maize and *Tripsacum* have been independently reducing their gene content after their shared ancient tetraploidy event. About 60% of maize homeolog pairs have been reduced to singletons in the maize genome so far (Schnable et al., 2011). Given enough time, more than 90% of gene copies are eventually lost in eukaryotic genomes after whole genome duplication (Sankoff et al, 2010). Gene loss happens slowly; the half-life for duplicated genes in the *Arabidopsis thaliana* genome is 17.3 million years (Lynch and Conery, 2003). Thus, the maize and *Tripsacum* genomes are still shedding duplicated genes. Understanding which genes are under strong purifying selection and resistant to pseudogenization in *Tripsacum* and maize would greatly aid crop improvement efforts, however, the maize and *Tripsacum* transcriptomes have yet to be compared to identify a common gene set that is resistant to pseudogenization and gene loss since their shared polyploidy event.

Genes that are resistant to pseudogenization in maize and *Tripsacum* are predicted to be located near recombination hotspots. High recombination rates are associated with purifying selection and the removal of deleterious alleles (Rodgers-Melnick, 2015; Tiley and Burleigh, 2015). Recombination unlinks beneficial variants from deleterious mutations, allowing purifying selection to act more efficiently. Recombination also strongly influences nucleotide composition through GC-biased gene conversion. When recombination occurs at heterozygous sites, a heteroduplex forms. The mismatch repair machinery often favors the G/C allele over the A/T allele, leading to GC-biased gene conversion. GC-biased gene conversion increases the GC content in all three codon positions as well as introns and intergenic recombination hotspots. GC-biased gene conversion plays a major role in shaping GC content in maize, rice, and other plant genomes (Rodgers-Melnick, 2015; Muyle et al., 2011; Serres-Giardi, 2012). Although base composition is shaped by other evolutionary forces such as mutational bias and selection on codon usage, GC-biased gene conversion has emerged as the strongest force affecting base composition (Clement et al., 2017). Thus, genes resistant to gene loss in maize and *Tripsacum* are predicted to have higher recombination rates and higher GC content than other genes in the genome, although this has yet to be tested.

Here, we present high-quality *de novo* transcriptome assemblies for two *Tripsacum* species: *Tripsacum dactyloides* and *Tripsacum floridanum* Porter ex Vasey. We hypothesize that a subset of genes shared by *Tripsacum* and maize are resistant to gene loss in both clades. Genes supported by experimental protein evidence in maize are good candidates for genes that are resistant to pseudogenization and gene loss in maize and *Tripsacum*. Functional genes under purifying selection produce protein, while pseudogenes cannot produce protein due to disabling mutations affecting the coding sequence (Xiao et al., 2016). A recent study by Walley et al. (2016) found that less than half of maize RefGen_v4 genes (15,160 out of 39,324) are detected in the proteome, which was constructed using electrospray ionization tandem mass spectrometry (Walley et al., 2016). Genes with detected protein tend to be syntenically conserved with sorghum, indicating that they are under high selective pressure (Walley et al., 2016). We tested whether maize genes with protein evidence and their *Tripsacum* homologs have higher GC content, higher expression, and more tightly conserved expression levels than putatively untranslated maize genes and their *Tripsacum* homologs. We also tested whether expression dominance in homeologous gene pairs was conserved across maize and *Tripsacum*.

## Materials and methods

### Plant material and tissue collection

Two mature *Tripsacum* individuals were used for this study. The first individual was from a *Tripsacum dactyloides* cultivar called “Pete” (PI 421612), which is a composite of 70 accessions native to Oklahoma and Kansas. Pete was developed through open-pollination and combine-harvesting over three generations. The rootstock for the Pete individual was acquired from the Tallgrass Prairie Center in Cedar Falls, Iowa in 2010 and given the internal identifier “IA_Pete_1”, passport #77. The second individual used for this study was a *Tripsacum floridanum* field collection obtained near Navy Wells, Florida in 2005. It was given the internal identifier “FL_05_15_1”, passport #8.

mRNA was harvested from the *Tripsacum dactyloides* individual and *Tripsacum floridanum* individual. These were mature potted plants that have been clonally propagated in the Cornell University greenhouses set at 22-24 °C. During this period, they have continually flowered and produced vegetative biomass. The *Tripsacum floridanum* tissue types collected for RNA extraction were mature roots, crown tissue (hardened green tissue located at the very base of the leaf bundles, just above the proaxis), whole mature leaves, and an inflorescence from three stages: pre-silking, post-silking, and post-anthesis. The same tissue types were collected for *Tripsacum dactyloides* except the post-silking and post-anthesis inflorescences. The *Tripsacum dactyloides* individual only had one inflorescence at the time of tissue collection. Tissue was harvested in the greenhouse between 2-3 pm and frozen immediately in liquid nitrogen.

### RNA extraction, library preparation, and sequencing

*Tripsacum* tissue was ground in liquid nitrogen using a mortar and pestle. RNA was extracted using the Direct-zol RNA miniprep kit with DNase I digestion. The *Tripsacum* RNA samples and one maize B73 RNA sample were used for library construction. The B73 RNA originated from mature adult leaf grown in a field in Aurora, NY collected during the day. Strand-specific cDNA libraries were constructed using the Lexogen SENSE mRNA-Seq library prep kit for Illumina V2. The RTL Reverse Transcription and Ligation Mix was used in the amounts suggested by Lexogen to achieve a mean insert size of 443 bp. Libraries were quantified using the KAPA library quantification kit for Illumina platforms. All libraries were multiplexed, and the pool was run on two separate lanes of the NextSeq 500 to generate 2×150 bp reads.

### Read processing for quality control

The following quality control processing steps were performed on the raw reads using Trimmomatic (version 0.32). Adapters listed in Supplemental Table S1 were removed with eight maximum allowed seed mismatches, a palindrome clip threshold of 30, a simple clip threshold of 11, and a minimum adapter length of one. Both reads were kept if they were exact reverse complements of each other. Reads were trimmed if the average Phred score in a five base sliding window fell below 20. Bases were trimmed from the beginning and end of a read if their Phred score was less than 35 or 20, respectively. All reads shorter than 36 bp were removed from the dataset. To remove contaminating rRNA, these quality-trimmed reads were aligned with Bowtie1 against the MIPS repetitive sequence element database (Nussbaumer et al., 2013). All reads that aligned to repeats were removed from the dataset. A final effort to remove contaminating Illumina adapter sequences involved aligning Illumina adapter sequence queries (Supplemental Table S1) to a Blast database of cleaned reads. All reads that aligned to adapter sequences were removed from the dataset (about 1,400 per library). Finally, to remove inaccurate bases from non-specific binding of the Lexogen primers during library construction, Trimmomatic clipped 5 bp and 3 bp from the beginning of the forward and reverse read respectively.

### *de novo* assembly and filtering of the transcriptomes

After cleaning the reads, all tissue-specific libraries were combined within each species for *de novo* transcriptome assembly. Trinity (version 2.1.1) assembled the transcriptomes using a minimum kmer coverage of two. To determine if short transcripts should be removed from the transcriptome assemblies, *Tripsacum* transcripts were aligned to maize B73 protein sequences (RefGen_v3) using Blastx and an e-value cut-off of 1×10^−20^. Transcripts shorter than 500 bp were removed from both assemblies.

Blobtools (Kumar et al., 2013) was used to remove transcripts from the assembly that originated from microbial or fungal species present in the environment and microbiome. In the Blobtools pipeline, cleaned reads were mapped back to the transcriptome assembly of their appropriate species using Bowtie2 (Langmead and Salzberg, 2012) with the --very-fast-local parameter. Transcript sequences from both species were used as a queries against the Blast nt database to determine the species of the best Blast hit. An e-value cutoff of 1e^−5^ was used for the Blast alignments. The blobology tool was used to plot each transcript by expression, GC content, and phylum. All transcripts that did not have a best Blast hit in Streptophyta were removed from the assembly.

### Evaluating the quality of the transcriptome assemblies

To measure how much of the input read space Trinity used in its original assemblies before filtering out low quality and contaminating transcripts, cleaned reads were mapped back to the transcriptome assemblies using Bowtie2. Bowtie2 was used with a maximum insert size of 1000 bp and the ‘--very-sensitive’ parameter. To calculate the L50 statistic for each expression level, it was first necessary to quantify transcript abundance within each library. Reads from each tissue-specific library were mapped back to the filtered transcriptomes using the Trinity script ‘align_and_estimate_abundance.pl’ that calls Bowtie2 and eXpress. A maximum insertion size of 1000 bp was used. For each species, transcripts were divided into ten different expression bins based on the TMM-normalized transcripts per million (TPM) statistic reported by eXpress. There were an equal number of transcripts in each expression bin. The L50 statistic was calculated for the transcripts in each expression bin.The definition of L50 is that half of all the assembled bases in each bin exist in transcripts at least as long as the L50 length.

The transcript assemblies were assessed with BUSCO version 2 (Simao et al., 2015). All assembled transcripts within each *Tripsacum* species were submitted as queries against the core eukaryotic gene set for flowering plants (embryophyta_odb9). Duplicate BUSCO hits were removed if they were homologs for the same maize gene. If they had no maize homolog, duplicate BUSCO hits were removed if they were grouped within the same Trinity gene cluster.

### Finding maize homologs of *Tripsacum* transcripts

*Tripsacum dactyloides* and *Tripsacum floridanum* transcripts were aligned to the B73 RefGen_v4 genome using PASA (Haas et al., 2003). PASA was run using gmap as the aligner. PASA was run with the following requirements for a valid alignment: a maximum intron length of 20,000 bp, a minimum average percent identity of 80%, a minimum aligned length of 70%, and no base pairs flanking the splice junctions were required to match perfectly. PASA reported only one valid alignment per transcript.

A *Tripsacum* transcript was considered a homolog of a maize gene if: 1) part of the transcript alignment fell within the boundaries of the maize gene and aligned to the same genomic strand as the maize gene, 2) the best Blastx hit for the *Tripsacum* transcript was a protein for that maize gene (RefGen_v3), and 3) the *Tripsacum* transcript alignment covered at least 95% of the length of the best Blastx maize protein hit (RefGen_v3). A single maize transcript from each maize gene homolog was considered the best homolog for a *Tripsacum* transcript if: 1) the maize RefGen_v4 transcript aligned to the same maize RefGen_v3 protein as the fully-assembled *Tripsacum* transcript, 2) the maize RefGen_v4 transcript and maize RefGen_v3 protein were expressed from the same gene according to the gene ID history between version 3 and version 4 of the maize genome (using the file available at the time: maize.v3TOv4.geneIDhistory.txt), and 3) the RefGen_v4 transcript had the longest alignment length to the RefGen_v3 protein out of any other transcript from the RefGen_v4 gene. The RefGen_v3 annotations were used because the RefGen_v4 annotations were under revision at the time. The transcript homolog pairs that were between 500 bp and 10,000 bp long were analyzed for GC content using in-house scripts.

### Characterizing non-maize transcripts

*Tripsacum* transcripts that did not align to the RefGen_v4 genome using PASA were aligned to maize B73 RefGen_v3 cDNA using Blastn. A discontiguous megablast parameter and an e-value threshold of 1e-20 was used. The expression values of transcripts were obtained from the previous eXpress run. Transcripts with an FPKM greater than one were used as a Blastx query against the NCBI non-redundant protein database. The reading frame with the best Blast hit was considered the open reading frame. Reverse open reading frames were not used because the transcripts were assembled in a strand-specific manner. The EMBOSS transeq (Rice et al., 2000) software was used to get translations in the open reading frame. Conserved protein domains within these translations were identified using the HMMER (version 3.1b2) hmmscan command. The protein sequences were used as a query against the Pfam A database with an inclusion threshold of 0.01 (Eddy, 2009; Finn et al., 2016).

### B73 vs. *Tripsacum* expression

The following quality control processing steps were performed on the raw B73 reads using Trimmomatic (version 0.36). Adapters listed in Supplemental Table S1 were removed with eight maximum allowed seed mismatches, a palindrome clip threshold of 30, a simple clip threshold of 11, and a minimum adapter length of one. Both reads were kept if they were exact reverse complements of each other. All reads shorter than 36 bp were removed from the dataset. To remove inaccurate bases from non-specific binding of the Lexogen primers during library construction, Trimmomatic clipped 9 bp and 6 bp from the beginning of the forward and reverse read respectively. Reads were trimmed if the average Phred score in a five base sliding window fell below 20. Bases were trimmed from the end of a read if their Phred score was less than 20.

Mature adult leaf libraries from B73, *Tripsacum dactyloides*, and *Tripsacum floridanum* were aligned to the B73 genome (version RefGen_v4) using the STAR aligner (Dobin et al., 2012, version 2.5.2b). Read alignments were considered valid if they were unique and less than 6% of the read was mismatches. The RefGen_v4.34 annotations were during the STAR alignments. Each gene in the GTF file had its 3’ boundary extended by 500 bp. The -- sjdbOverhang parameter was used with a value of 89. Cufflinks2 (version 2.2.1) was then used to estimate transcript abundance. The intron size was limited to 20 − 80,000 bp. Gene FPKM values from Cufflinks2 were used to compare *Tripsacum* and maize expression. Genes with FPKM values lower than 0.01 in either species were not used for the comparison. For the homeolog expression analysis, the 1,750 homeologous pairs identified by Schnable et al. (2011) were used. Their RefGen_v4 names were identified using the maizeGDB conversion file v3_v4_xref.txt.

## Results

### *de novo* transcriptome assembly

Transcriptomes were assembled from root, leaf, crown, and inflorescence tissue from two species, *Tripsacum dactyloides* and *Tripsacum floridanum*, using Trinity (Grabherr et al., 2011). Transcripts shorter than 500 bp were filtered out of the assemblies because they were mostly fragmented and did not cover the full length of the nearest B73 maize homolog (Supplemental Figure S1). Contaminating transcripts from organisms such as fungi and insects in the environment or microbiome were removed using Blobtools scripts (Supplemental Figure S2) (Kumar et al., 2013).

In the final transcriptome assemblies, Trinity grouped the transcripts into 50,411 *Tripsacum dactyloides* genes and 64,422 *Tripsacum floridanum* genes (Table 1), which is considerably more than the 39,656 genes in the filtered RefGen_v3 maize gene set (Law et al., 2015). Trinity may have overestimated gene number if it misidentified allelic transcripts as transcripts originating from different genes. The sequenced individuals are highly heterozygous, so allelic transcripts were probably assembled for many genes. We do not attempt to collapse allelic transcripts at the gene level because our ultimate purpose was to identify homologous pairs at the transcript level in maize and *Tripsacum*. The final assemblies contain 131,952 *Tripsacum dactyloides* transcripts and 155,705 *Tripsacum floridanum* transcripts, respectively (Table 1).

**Table 1:** *de novo* transcriptome assembly statistics for *Tripsacum dactyloides* and *Tripsacum floridanum*

| Assembly statistic | *Tripsacum dactyloides* | *Tripsacum floridanum* |
| --- | --- | --- |
| Amount of High Quality Sequence Used For Assembly | 46 Gbp | 79 Gbp |
| Estimated Coverage of *Tripsacum* Transcriptome | 479X | 810X |
| Reads Mapped to Unfiltered Transcriptome Assembly | 93% | 93% |
| Transcript L50 | 1,442 bp | 1,452 bp |
| Median Transcript Length | 981 bp | 956 bp |
| Mean Transcript Length | 1,230 bp | 1,233 bp |
| Number of Trinity-defined Genes | 50,411 | 64,422 |
| Number of Trinity-defined Transcripts | 131,952 | 155,705 |
| Completely-assembled BUSCO genes | 83.8% | 85.5% |

### The transcriptome assemblies are high quality

Ideally, the average transcript length of a *de novo* transcriptome assembly should be similar to the average transcript length in the most closely related species. The mean transcript lengths were 1,230 bp for *Tripsacum dactyloides* and 1,233 bp for *Tripsacum floridanum*, while the mean transcript length is 1,541 bp in maize RefGen_v3 annotations (Law et al., 2015). The L50 was 1,442 bp and 1,452 bp for the *Tripsacum dactyloides* and *Tripsacum floridanum* assemblies, respectively. The L50 statistic defines the transcript length for which half of all assembled bases exist in transcripts longer than the L50 length. More highly expressed transcripts have longer L50 lengths, probably because higher read coverage depth enables a more complete assembly (Supplemental Figure S3).

The Benchmarking Universal Single-Copy Ortholog (BUSCO) gene set for land plants was used to assess the completeness of the two assemblies (Simao et al., 2015). Of all plant BUSCO genes, 83.8% and 85.5% were completely assembled in *Tripsacum dactyloides* and *Tripsacum floridanum*, respectively (Supplemental Figure S4). These data indicate that a large majority of genes in the genome are represented in the two assemblies.

### Identification of *Tripsacum* and maize homologs

Full-length *Tripsacum* transcripts, meaning transcripts that aligned to 95% of the length of its nearest maize protein homolog (RefGen_v3), were aligned to the maize inbred B73 genome (RefGen_v4) to identify their nearest maize gene homolog. Alignments were required to have at least 80% identity between maize and *Tripsacum* and at least 70% of the *Tripsacum* transcript length aligning to the genome. The average percent identity across all valid transcript alignments was 91.98% in *Tripsacum dactyloides* and 91.49% in *Tripsacum floridanum*. There were 6,982 homologous transcript pairs between *Tripsacum dactyloides* and maize, and there were 6,368 homologous transcript pairs between *Tripsacum floridanum* and maize. There were 1,737 *Tripsacum dactyloides* transcripts and 2,303 *Tripsacum floridanum* transcripts that did not align to the B73 genome or B73 cDNA and had an expression level greater than one FPKM in at least one tissue. These *Tripsacum* transcripts that are absent from the B73 maize genome are listed in Supplemental Data 1-2, along with their best Blast hits from the NCBI non-redundant protein database and their conserved Pfam domains. These *Tripsacum-specific* genes contain a broad variety of conserved protein domains and a diverse set of functions, and many are putative transcription factors.

### Proteome transcripts have higher GC content than non-proteome transcripts in maize and *Tripsacum*

We hypothesized that protein-encoding maize transcripts and their *Tripsacum* homologs (hereafter called “proteome pairs”) have maintained higher GC content than putatively untranslated maize transcripts and their *Tripsacum* homologs (hereafter called “non-proteome pairs”). The proteome study by Walley et al. (2016) was used to determine whether a maize transcript encoded a protein or not. Average GC content was calculated along the length of maize and *Tripsacum* homologous transcripts at all coding and non-coding positions. GC content peaks at the 5’ end of transcripts and declines toward the 3’-end in maize, *Tripsacum dactyloides*, and *Tripsacum floridanum* (Figure 1a,b). Other studies have also observed high GC content at the 5’-end of genes and a declining GC gradient over the length of the gene in monocots and dicots (Serres-Giardi, 2012; Wong, 2002). Recombination hotspots in monocots and dicots tend to be located at the 5’-end of genes, and to a lesser extent, at the 3’-end of genes (Hellsten, 2013; Choi, 2013; Singh, 2016). Thus, the observed declining GC gradient over transcript length is probably due to GC-biased gene conversion. Proteome pairs had about a 1-2% higher GC content peak at the 5’-end of transcripts than non-proteome pairs (Figure 1a-c). These results suggest that the regions of the genome where proteome pairs reside have higher recombination rates than the regions of the genome where non-proteome pairs reside.

**Figure 1:**
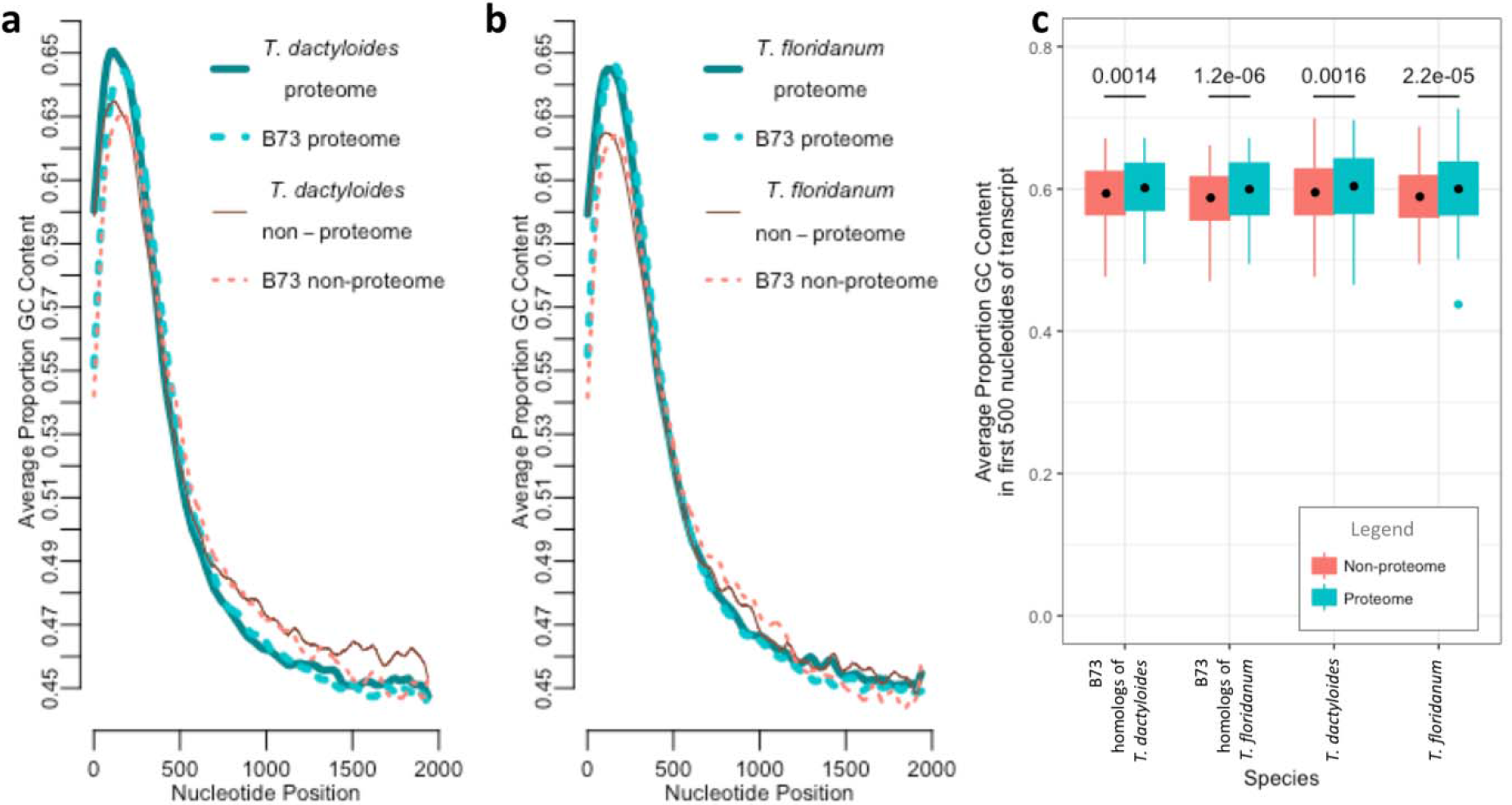
Average GC content in fully-assembled *Tripsacum* transcripts and their maize homologs. **a,b**) LOWESS curves of GC content averaged across transcripts at each nucleotide position. Homologous transcript pairs for maize and *T. dactyloides* (**a**) or maize and *T. floridanum* (**b**) were split into two subsets: pairs where the maize homolog is detected in the proteome (Walley et al., 2016), and pairs where the maize homolog is not detected in the proteome. **c**) Boxplot of the average proportion GC content in the first 500 nucleotides of transcripts. P-values are shown for the Student’s two-sided t-test with the Welch approximation. The turquoise point is an outlier, and the black points represent the mean of the average proportion GC content across the first 500 nucleotides.

### Proteome genes have higher, more conserved expression than non-proteome genes in maize and *Tripsacum*

If proteome genes are more resistant to gene loss in maize and *Tripsacum*, they will have higher expression than non-proteome genes because highly expressed genes are more resistant to genome fractionation than lowly expressed genes in maize (Schnable et al., 2011; Zhao et al. 2017). Gene expression levels were estimated using maize and *Tripsacum* RNA-seq reads that mapped uniquely to the maize genome. The percentage of reads that mapped uniquely for each species was 60.81% in *Tripsacum dactyloides*, 53.86% in *Tripsacum floridanum*, and 81.72% in maize. The average alignment mismatch rate was 2.58% in *Tripsacum dactyloides*, 2.57% in *Tripsacum floridanum*, and 0.19% in maize. Homologous genes with FPKM values over 0.01 in both species are plotted in Figure 2. A two-sample t-test with the Welch approximation was performed to test whether protein-encoding maize genes and their *Tripsacum* homologs (“proteome pairs”) were expressed at different levels than non-proteome maize genes and their *Tripsacum* homologs (“non-proteome pairs”). Proteome pairs had significantly higher mean log-transformed FPKM values than non-proteome pairs in maize (t(22176) = 39.7, p < 2.2 × 10^−16^), *Tripsacum dactyloides* (t(22728) = 54.3, p < 2.2 × 10^−16^), and *Tripsacum floridanum* (t(22332) = 50.6, p < 2.2 × 10^−16^) (Figure 2). The average un-transformed maize expression level was 30.0 FPKM for proteome genes and 13.3 FPKM for non-proteome genes. The average un-transformed *Tripsacum dactyloides* expression level was 36.5 FPKM for homologs of proteome genes and 14.4 FPKM for homologs of non-proteome genes. The average un-transformed *Tripsacum floridanum* expression level was 39.0 FPKM for homologs of proteome genes and 14.8 FPKM for homologs of non-proteome genes. Thus, proteome maize genes and their *Tripsacum* homologs are more highly expressed than non-proteome maize genes and their *Tripsacum* homologs.

**Figure 2:**
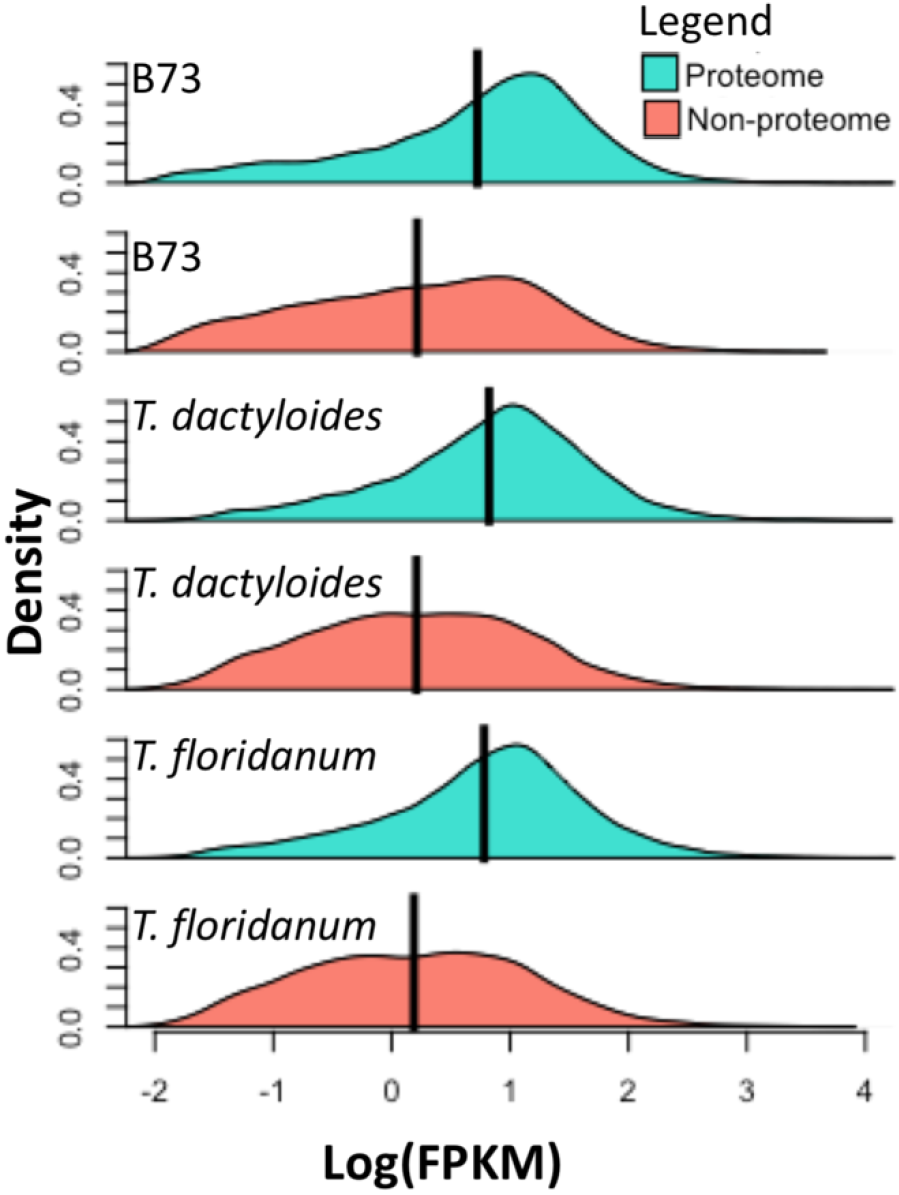
Probability density plots of gene expression levels in maize inbred B73, *Tripsacum dactyloides*, and *Tripsacum floridanum*. The black line is the mean log-transformed expression level. mRNA-seq reads from mature adult leaves were aligned to the B73 genome (RefGen_v4) using Cufflinks2. Maize genes were divided into two subsets: genes translated into the maize proteome (Walley et al., 2016), and genes not detected in the proteome. The log-transformed expression of genes with an FPKM > 0.01 are plotted.

We hypothesized that proteome pairs had more conserved gene expression levels than non-proteome pairs. Simple linear regressions were performed to predict *Tripsacum dactyloides* (Figure 3a-c) and *Tripsacum floridanum* (Figure 3d-f) gene expression based on maize gene expression. The expression levels of proteome maize genes explain 42% of the variance in *Tripsacum dactyloides* homolog expression (Figure 3b) and 49% of the variance in *Tripsacum floridanum* homolog expression (Figure 3e). The expression levels of non-proteome maize genes explain less variance - only 29% of the variance in *Tripsacum dactyloides* homolog expression (Figure 3c) and 37% of the variance in *Tripsacum floridanum* homolog expression (Figure 3f). These data indicate that translated maize genes have more tightly conserved expression with *Tripsacum* than non-proteome maize genes. Gene expression levels are even more tightly conserved within the *Tripsacum* genus. The expression levels of *Tripsacum floridanum* homologs of maize proteome genes explain 78% of the expression levels of *Tripsacum dactyloides* homologs of maize proteome genes (Figure 3h). Interestingly, the expression levels of non-proteome homologs are also tightly conserved within *Tripsacum* because the regression model explains 72% of the variance (Figure 3i). This indicates that the expression levels of non-proteome homologs have not diverged much within the *Tripsacum* genus but have become more highly diverged between the *Tripsacum* and maize genera.

**Figure 3:**
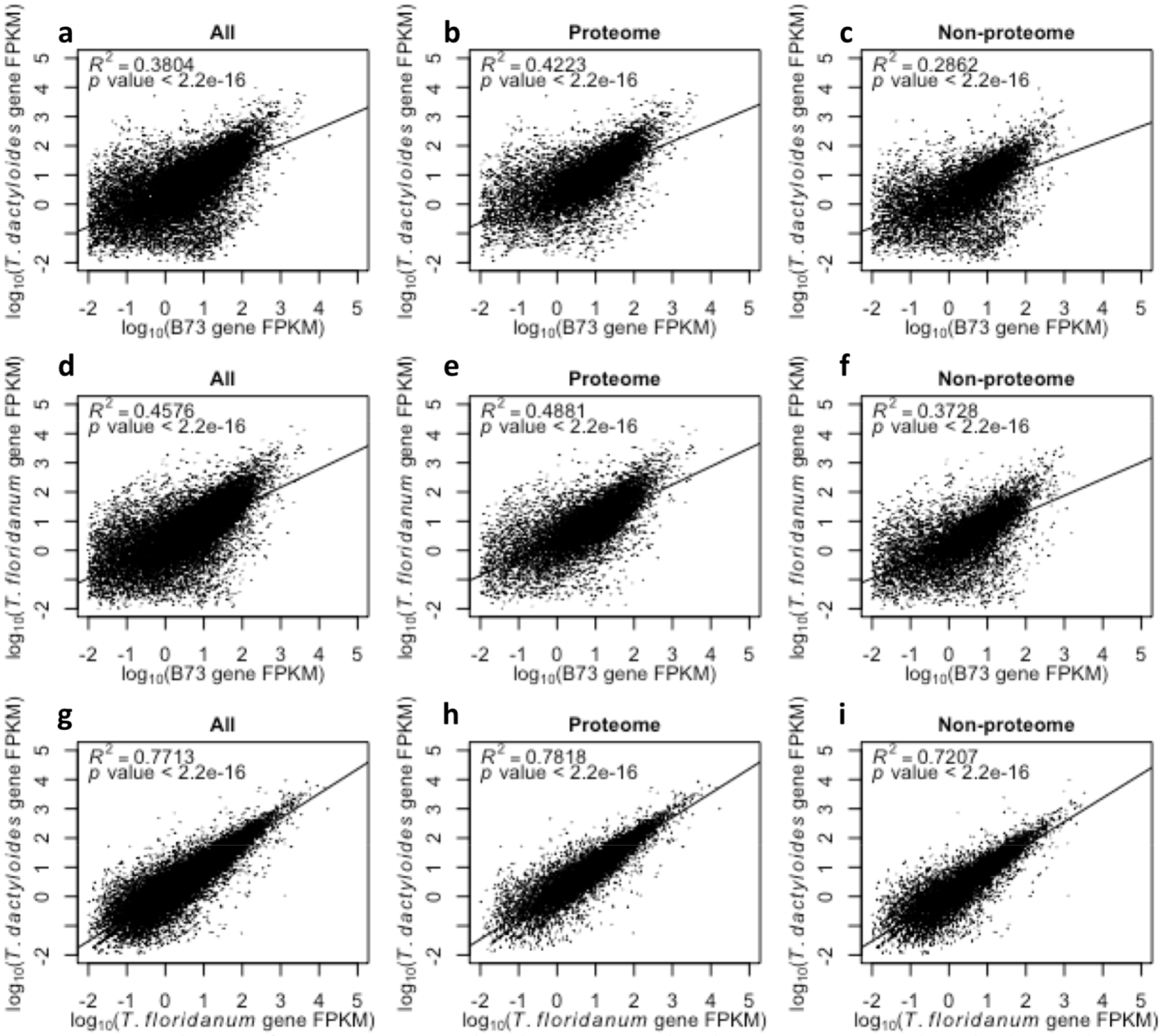
Correlation of gene expression levels in maize inbred B73, *Tripsacum dactyloides*, and *Tripsacum floridanum*. RNA-seq reads from mature adult leaves were aligned to the B73 genome (RefGen_v4) using Cufflinks2. Maize genes were divided into two subsets: genes translated into the maize proteome (Walley et al., 2016), and genes not detected in the proteome. The log-transformed expression of genes with an FPKM > 0.01 were plotted in B73 vs. *Tripsacum dactyloides* (**a-c**), B73 vs. *Tripsacum floridanum* (**d-f**), and *Tripsacum floridanum* vs. *Tripsacum dactyloides* (**g-i**). Adjusted *R*^2^ values are shown.

**Figure 4:**
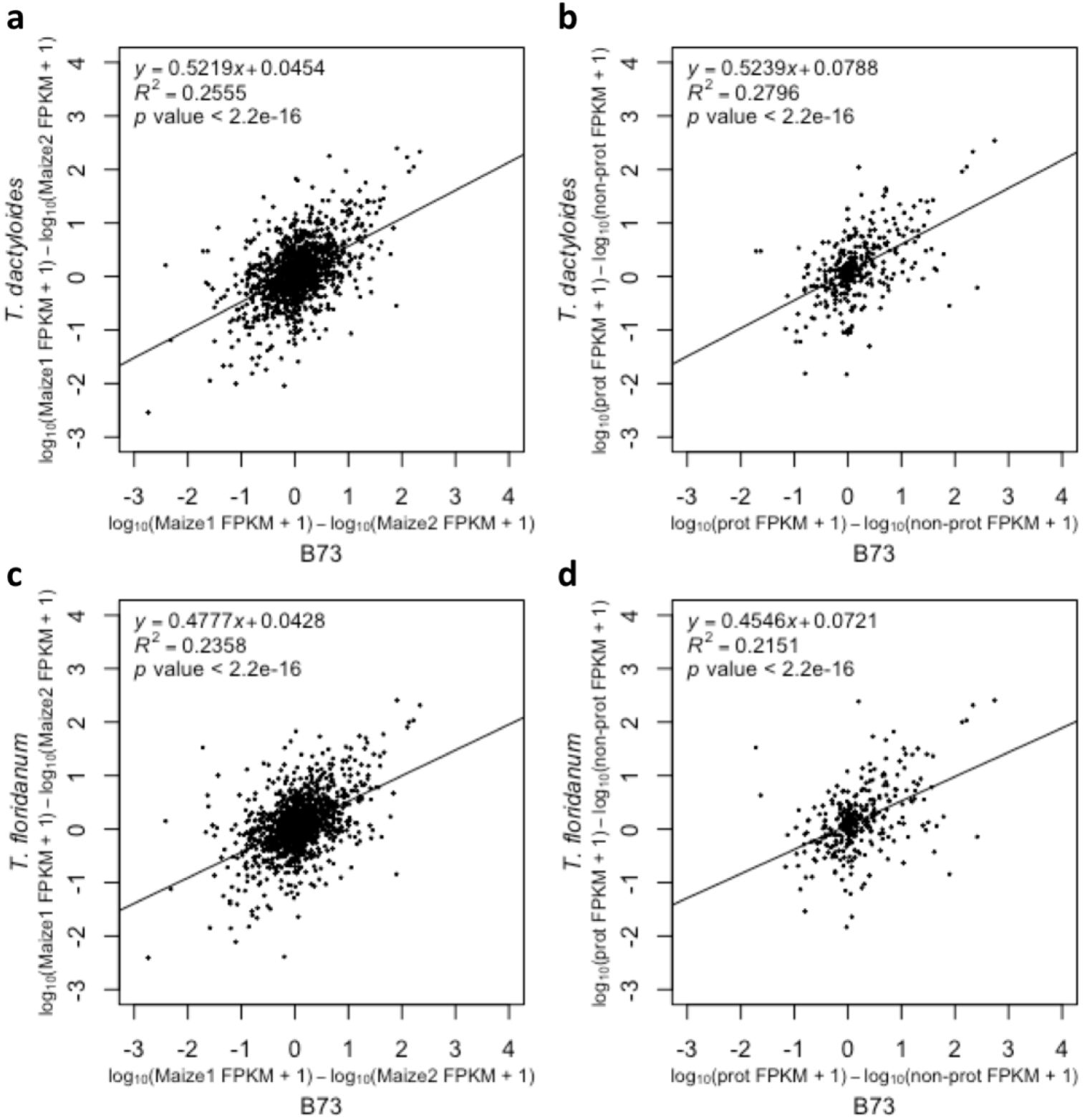
Correlation of homeolog expression differences in maize inbred B73 and *Tripsacum dactyloides* (a,b) and maize inbred B73 and *Tripsacum floridanum* (c,d). Difference in log-transformed FPKM values between maize subgenome 1 homeolog and maize subgenome 2 homeolog is plotted for all 1,625 homeolog pairs (a, c). Difference in log-transformed FPKM values between the homeolog detected in the proteome and the homeolog not detected in the proteome is plotted for 367 homeolog pairs (b, d). A unit of 1 FPKM was added to all gene expression values to aid in log transformation of unexpressed genes. Adjusted *R*^2^ values are shown.

### The same homeolog tends to have higher expression in maize and *Tripsacum* in reciprocally retained homeolog pairs

If genome fractionation is occurring in a similar fashion in maize and *Tripsacum*, then the same member of a homeolog pair will exhibit higher expression than other member in both clades. There are 1,625 homeolog pairs identified by Schnable et al. (2011) that have homologs in our *Tripsacum dactyloides* and *Tripsacum floridanum* transcriptomes. Each homeolog pair consists of a homeolog from the maize 1 subgenome and a homeolog from the maize 2 subgenome. The difference in maize 1 homeolog expression and maize 2 homeolog expression are significantly correlated between maize and *Tripsacum* (Figure 4a,c). Maize homeolog expression differences explain 26% of *Tripsacum dactyloides* homeolog expression differences (Figure 4a) and 24% of *Tripsacum floridanum* homeolog expression differences (Figure 4c). Of the 367 maize homeolog pairs that had one proteome gene and one gene that was not detected in the maize proteome, the homeolog expression differences in maize explained 28% of the variation in expression differences in *Tripsacum dactyloides* (Figure 4b) and 22% of the variation in expression differences in *Tripsacum floridanum* (Figure 4d). These data indicate that the same homeolog tends to show expression dominance in maize and *Tripsacum*, but it isn’t always necessarily the maize 1 homeolog or the proteome homeolog.

Expression levels often differ between members of homeologous gene pairs. In soybean, about half of homeologous gene pairs exhibited transcriptional divergence between members in at least one tissue (Roulin et al., 2013). In a natural cotton allopolyploid (*Gossypium L.*), about 40% of 49 homeologous gene pairs showed transcriptional bias toward one homeolog or the other (Chaudhary et al., 2009). In maize, 98% of 3,228 homeologous gene pairs exhibited two-fold transcriptional divergence between members across multiple tissues (Pophaly et al., 2015). The more highly expressed maize homeolog had significantly lower *Πn/Πs* ratios, and thus were under stronger purifying selection, than the more lowly expressed homeolog (Pophaly et al. 2015). In this study, we do not test for differential expression of genes within homeologous pairs, but we do see that the polarity in homeolog expression is conserved between maize and *Tripsacum*.

## Discussion

We tested whether the gene complements of maize and *Tripsacum* are reducing in size in a similar fashion following their shared ancient tetraploidy event. Maize genes with protein evidence were promising candidates for genes resistant to pseudogenization in both clades. Maize genes that produce detectable proteins tend to be syntenic (Walley et al., 2016), and syntenic genes tend to be more highly conserved (Schnable, 2015). With this study, we show that the maize proteome genes also have higher GC content and higher expression than non-proteome maize genes (Figures 1,2). Furthermore, *Tripsacum* homologs of maize proteome genes also have higher GC content and gene expression than *Tripsacum* homologs of maize non-proteome genes (Figures 1,2). Expression patterns are more tightly conserved across maize and *Tripsacum* for proteome pairs than non-proteome pairs (Figure 3). The same gene in homeologous pairs tends to show expression dominance in both maize and *Tripsacum* (Figure 4). Together, these data indicate that the maize and *Tripsacum* gene sets are evolving in parallel after whole genome duplication.

In this study, GC content served as an indirect measure of recombination rate because exonic and intronic GC content is positively correlated with recombination rate due to GC-biased gene conversion (Muyle et al. 2011). GC content in exonic positions peaked near the 5’-end of the gene and steeply declined toward the 3’-end. Because recombination hotspots in monocots and dicots tend to be located at the 5’-end of genes (Hellsten, 2013; Choi, 2013; Singh, 2016), this decline in GC content is likely due to GC-biased gene conversion. The higher the heterozygosity, the stronger the effect of GC-biased gene conversion (Muyle et al., 2011). GC-biased gene conversion likely has such a large effect because *Tripsacum* and maize populations are highly heterozygous. Thus we conclude that proteome pairs have a higher GC content, and by proxy a higher recombination rate, than non-proteome pairs.

One potential limitation of this study is that a single biological leaf replicate was used for the maize, *Tripsacum dactyloides*, and *Tripsacum floridanum* expression analyses. Single replicates were taken from root, leaf, crown, and inflorescence tissue with the purpose of sampling transcript diversity to assemble the *Tripsacum* transcriptomes. When the leaf RNA-seq data was later used for interspecific comparisons of expression, the same patterns in gene expression between maize and *Tripsacum dactyloides* are also evident when comparing maize and *Tripsacum floridanum. Tripsacum dactyloides* and *Tripsacum floridanum* behave as independent replicates for the *Tripsacum* genus. Both replicates indicate that maize proteome gene expression is more tightly with *Tripsacum* than maize non-proteome gene expression. Furthermore, each homeolog pair represents a separate opportunity for expression divergence to occur. The same homeolog tended to have higher expression in each of the three species.

While the exact nature of maize genes lacking protein evidence needs further study, this gene set may be enriched for pseudogenes or genes on the evolutionary path toward pseudogenization. Pseudogenes can form after a polyploidy event when there is a lack of selective pressure to maintain two copies, and one homeolog loses the ability to produce a functional protein (Xiao et al., 2016). Although the two homeologs have similar gene sequences after whole genome duplication, they differ in genomic location as large-scale chromosomal rearrangements occur. If one homeolog has a lower recombination rate, it is more likely to accumulate deleterious mutations. Mutations or small intra-exon deletions (Woodhouse et al., 2010) that disrupt the open reading frame can cause frame shifts or premature termination codons and may transform a gene into a pseudogene. Several studies have shown that pseudogenes can be actively transcribed. mRNA was detected for 12% of rice pseudogenes, which have significantly lower expression levels than their functional paralogs (Thibaud-Nissen et al., 2009). Out of 129 lineage-specific pseudogenization events in four Poaceae species, 61 pseudogenes are actively transcribed (Zhao et al. 2015). Zou et al. (2009) found that 17% and 23% of pseudogenes are expressed in rice and *Arabidopsis*, respectively, and they have similar or lower expression than functional genes. Pseudogenes may be mis-annotated as functional genes using *ab initio* gene finding software, which can alter gene structure in order to avoid frame shift mutations or premature stop codons (Thibaud-Nissen et al., 2009).

Despite a thorough sampling of 33 tissues, some proteins may be missing from the Walley et al. (2016) proteome if they accumulate in stress conditions or at certain times in the diurnal cycle that were not sampled. Some proteins may not have been detected due to limited sensitivity of the mass spectrometer. Thus, some genes we have been referring to as “non-proteome genes” may not be pseudogenes because they do in fact produce functional proteins. More experiments are needed to determine whether these genes without detectable protein are truly undergoing pseudogenization.

The expression analysis between 1,625 high-confidence homeolog pairs identified by Schnable et al. (2011) in maize and *Tripsacum* reveals parallel gene evolution. The same homeolog tends to have expression dominance in both maize and *Tripsacum*. The proteome homeolog does not always have higher expression than the non-proteome homeolog. Similarly, the maize 1 homeolog does not always have higher expression than the maize 2 homeolog. Likewise, it may be expression dominance itself that determines whether a homeolog is more likely to resist fractionation. To support this notion, expression dominance is better indicator than subgenome location for whether a homeolog is under strong purifying selection (Pophaly et al., 2015).

Natural selection is shaping the gene sets of maize and *Tripsacum* similarly after tetraploidy. Maize breeding programs could prioritize genes that are highly expressed, protein-encoding, and conserved across genera because they are likely to control plant phenotype. Because less than half of maize genes fit these criteria, it reduces the number of genes that breeders have to consider targeting. This study represents one application out of many possible applications for how the *de novo* transcriptome assemblies presented here can be used to provide insight into maize genome evolution.

## Acknowledgements

This material is based upon work by C.M.G. supported by the National Science Foundation Postdoctoral Research Fellowship in Biology under Grant No. 1523861. This material is also based upon work by K.A.K. supported by the National Science Foundation Graduate Research Fellowship under Grant No. DGE-1650441. This work was supported by the US Department of Agriculture-Agricultural Research Service.

## Author contributions

C.M.G. and E.S.B. designed research. C.M.G. performed research. C.M.G., K.A.K., and E.S.B. analyzed data. C.M.G. wrote the paper.

## Supplemental material

Supplemental material includes supplemental figures and tables is available online.

## Data deposition

The accession numbers for the transcriptome assemblies reported in this paper will be publicly available soon.

